# Maize centromeric chromatin scales with changes in genome size

**DOI:** 10.1101/2020.11.05.370262

**Authors:** Na Wang, Jianing Liu, William A. Ricci, Jonathan I. Gent, R. Kelly Dawe

**Author notes:** Corresponding author: R. Kelly Dawe, Department of Genetics, B414A Davison Life Sciences, University of Georgia, Athens, GA 30602.

## Abstract

Centromeres are defined by the location of Centromeric Histone H3 (CENP-A/CENH3) which interacts with DNA to define the locations and sizes of functional centromeres. An analysis of 26 maize genomes including 110 fully assembled centromeric regions revealed positive relationships between centromere size and genome size. These effects are independent of variation in the amounts of the major centromeric satellite sequence CentC. We also backcrossed known centromeres into two different lines with larger genomes and observed consistent increases in functional centromere sizes for multiple centromeres. Although changes in centromere size involve changes in bound CENH3, we could not mimic the effect by overexpressing CENH3 by threefold. Literature from other fields demonstrate that changes in genome size affect protein levels, organelle size and cell size. Our data demonstrate that centromere size is among these scalable features, and that multiple limiting factors together contribute to a stable centromere size equilibrium.

## Introduction

The histone variant known as CENP-A/CENH3 recruits a set of constitutive centromere proteins that in turn recruit the kinetochore proteins that interact with microtubules (Zhong *et al.* 2002; Kixmoeller *et al.* 2020; Mitra *et al.* 2020). When CENH3 binds to DNA, that sequence is referred to as centromeric DNA. How and where CENH3 binds is determined by a host of factors that differ among species. In human and mouse, centromere sequences can specifically promote the deposition of overlying centromere proteins (Aldrup-MacDonald *et al.* 2016; Iwata-Otsubo *et al.* 2017) although the fact that centromeres occasionally form on regions that lack canonical centromere repeats suggests that the underlying mechanism is epigenetic (Murillo-Pineda and Jansen 2020). This contrasts with maize and other plants where centromere sequences vary dramatically and the specification mechanisms are largely or entirely epigenetic (Oliveira and Torres 2018). Whether DNA sequence helps to guide the deposition of centromere proteins or not, centromere/kinetochore domains adopt predictable sizes and are stably propagated (Bodor *et al.* 2014; Gent *et al.* 2017).

Under the epigenetic model for centromere positioning, the location and size of the existing centromere is used as a template for the location of a newly replicated centromere (Mitra *et al.* 2020). However, we have observed extensive plasticity in the size and locations of centromeres that presumably represents both stochastic and physiological variation (Gent *et al.* 2015, 2017). The observed plasticity fits well with the proposal that centromeres are highly dynamic and their average size is determined in part by the concentration of kinetochore proteins (Bodor *et al.* 2014). An analysis of multiple grass species demonstrated that the sum of all kinetochore sizes in a cell scales linearly with genome size (Bennett *et al.* 1981; Zhang and Dawe 2012), suggesting that the amount of kinetochore proteins is at least partially dependent on genome size and cell volume (Zhang and Dawe 2012). As a test, maize chromosomes were introduced into the larger oat genomic background and centromere size measured by the amount of DNA occupied by CENH3 as interpreted by ChIP-seq. The maize centromeres increased in size by two-fold in the oat background as predicted (Wang *et al.* 2014). These results mirror a variety of studies showing that subcellular structures frequently scale with genome and cell size (Price *et al.* 1973; Gregory 2001; Cavalier-Smith 2005; Gillooly *et al.* 2015; Robinson *et al.* 2018).

As outlined by Marshall (Marshall 2016), cellular scaling could occur by several mechanisms. The simplest is the limiting precursor model, where the amount of a key component increases with cell size and directly contributes to the size of the structure of interest. In the case of centromeres, a likely candidate is CENH3/CENP-A. Prior data from *Drosophila* and human cell lines have shown that overexpression of CENP-A causes ectopic centromere formation in non-centromeric regions (Heun *et al.* 2006; Shrestha *et al.* 2017). Other likely limiting components are those involved in CENP-A deposition. CENP-A loading involves licensing factors such as Kinetochore Null 2 (KNL2) (Lermontova *et al.* 2013; Sandmann *et al.* 2017; Boudichevskaia *et al. 2019)*, specific chaperones (Sanchez-Pulido *et al.* 2009; Chen *et al.* 2014) and interactions with other kinetochore proteins such as Centromere Protein C (CENP-C) (Sandmann *et al.* 2017; French *et al.* 2017). CENH3/CENP-A, its licensing factors, chaperones, and other inner kinetochore proteins may directly or indirectly regulate centromere size either alone or in combination.

In the current work, we tested the idea that maize centromeres are scalable by analyzing recent genome assemblies of multiple inbreds, experimentally manipulating genome size using genetic crosses, and overexpressing CENH3. We find no consistent association between specific sequences and centromere size, and no change in centromere size after overexpressing *CENH3* by threefold. However, we found evidence of centromere scaling among inbreds that naturally vary in genome size and in lines with experimentally manipulated genome sizes. The data support the conclusion that centromere size is not controlled by DNA sequence or by CENH3 alone, but by a mass-action mechanism that is sensitive to cell volume and regulated by the concentration of multiple precursors.

## Results

### Limited impact of sequence on centromere location or size

In maize, the major centromere repeats are a tandem repeat known as CentC and an abundant class of Gypsy transposons known as Centromeric Retroelements (Wolfgruber *et al.* 2009). While both components are repetitive, they are diverse enough that many maize centromeres have been fully assembled, and a surprising number of short reads align to the assembled centromeres uniquely (Gent *et al.* 2012, 2015, 2017). For instance, seven B73 centromeres (2, 3, 4, 5, 8, 9 and 10) were assembled gaplessly in the recent B73-Ab10 assembly (Liu *et al.* 2020). This makes it possible to identify the sequence occupied by centromeric nucleosomes by aligning CENH3 ChIP-seq data from each inbred to the subset of assembled centromeres from that inbred. Functional centromere sizes can be estimated by identifying regions where the depth of ChIP-seq reads exceed an enrichment threshold, and enforcing a minimal peak size and maximal distance between peaks (Supplemental Figure 1 and Methods). Throughout this study, we only analyzed centromeres that were fully scaffolded where sequence gaps (if any) were of known size.

To assess natural centromere variation among a variety of maize lines, we measured centromere size in the 26 NAM founder inbreds (McMullen *et al.* 2009), for which high quality de novo genome assemblies have recently been completed (www.maizegdb.org/NAM_project). CENH3 ChIP-seq data are available for most of the NAM inbreds (Schneider *et al.* 2016). We carried out CENH3 ChIP-seq for eight of the NAM lines, either because the data were absent from the prior study, showed poor ChIP efficiencies, or because we wanted biological replicates prepared under the same conditions (Supplemental Table 1). Alignment of ChIP-seq data to the assemblies revealed that of the 260 centromeres present, 110 centromeres were fully scaffolded (Figure 1A). Of these, 88 were assembled gaplessly and 22 contained one or more gaps of known size.

**Figure 1.**
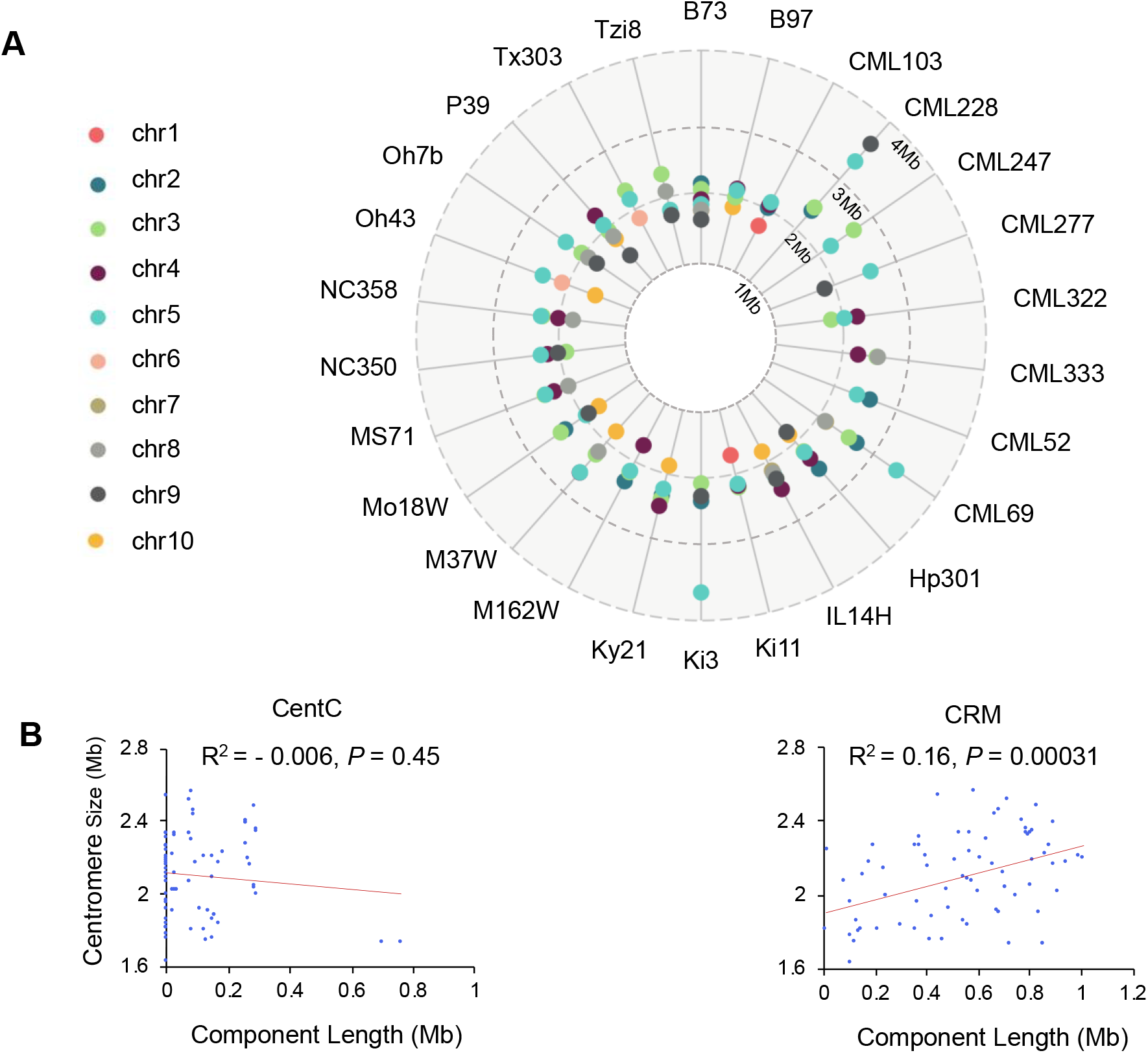
Centromeric repeats and their relationship to centromere size in the 26 NAM inbred lines. (A) The sizes of 110 fully assembled centromeres in each inbred line. (B) Linear regression analysis between centromere size and the abundance of CentC and CRM. Correlation coefficients and p-values are indicated in the graph.

In humans the alpha satellite actively recruits CENP-A and directly contributes to centromere size (Aldrup-MacDonald *et al.* 2016; Iwata-Otsubo *et al.* 2017). Our analysis of the CENH3-occupied regions of the 110 fully scaffolded centromeres revealed that larger centromeres do not necessarily have more CentC, although they do have more Centromeric Retroelements (CRM) (Figure 1B). The close relationship between CRM and centromeres appears to be a consequence of a specialized retroelement targeting mechanism (Schneider *et al.* 2016), but it remains possible that repeated CRM insertions could expand functional centromere size. Neither repeat, however, is found exclusively with functional centromeres: 59.5% of the CRM elements and 30.1% of the CentC repeats in the assembled reference genomes lie in flanking pericentromeric areas. The actual percentages may be smaller given that unknown amounts of each are present in the gaps of the 150 centromeres excluded from this analysis. Nonetheless, it is clear that large expanses of CRM and CentC can exist outside of functional centromeres.

### Positive correlation between centromere size and genome and chromosome sizes in NAM inbreds

The NAM founder inbreds were drawn from a wide genetic and demographic range, including tropical and northern lines as well as popcorn and sweet corn (McMullen *et al.* 2009). The genome sizes among the NAM founder inbreds vary from 2.09 to 2.50 Gb, where most of the differences in genome size can be attributed to differences in the amount of tandem repeat arrays within heterochromatic regions known as knobs (Chia *et al.* 2012).

We plotted the 110 measured centromere sizes against genome size and chromosome size. The data revealed a weak but significant positive correlation between centromere size and genome size (Figure 2A), supporting a prior cross-species study that came to the same conclusion (Zhang and Dawe 2012). Although we and others have speculated that the chromosomes within a species each have centromeres of similar size (Moens 1979; Zhang and Dawe 2012), our diverse collection of assembled centromeres revealed that larger chromosomes accommodate larger centromeres (Figure 2B), consistent with an earlier report from human cells showing a minor correlation between chromosome size and the number of attached microtubules (McEwen *et al*. 1998).

**Figure 2.**
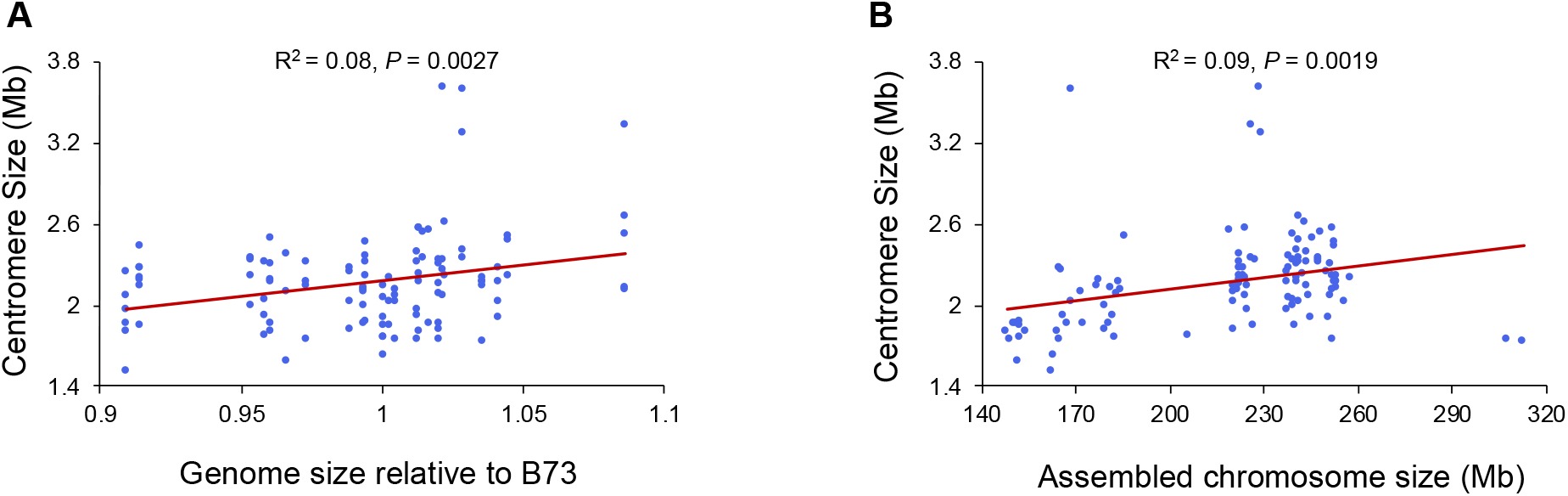
Correlation of centromere size with genome size and chromosome size. The graphs show data from 110 fully assembled centromeres in the 26 NAM genomes. (A) Linear regression analysis between genome size and centromere size. (B) Linear regression analysis between chromosome size and centromere size. Correlation coefficients and p-values are indicated in the graph.

### Centromeres expand when introduced into larger genomes

The only empirical data supporting a correlation between centromere size and genome size comes from an unnatural wide cross between maize and oat (Wang *et al.* 2014). We sought to confirm these results using natural crosses between the B73 inbred and two different genetic backgrounds: a maize landrace from Oaxaca Mexico with a genome about 1.3 times the size of the B73 genome, and the intercrossing species *Zea luxurians* with a genome size about 1.6 times the size of B73. The genomes of these two accessions are larger primarily because they contain more heterochromatic knob repeats (Bilinski *et al. 2018)*, although *Zea luxurians* also contains a larger proportion of retroelements (Tenaillon *et al.* 2011). Figure 3 shows the basic crossing schemes. We first crossed B73 with either Oaxaca or *Zea luxurians* to create F1s, which were self-crossed to create F2s or crossed again to the larger-genome parent to obtain BC1 lines. The BC1 lines were then self-crossed to create BC1F2 lines segregating for B73 centromeres. The genome sizes were measured for each cross using flow cytometry (Figure 3B and Supplemental Table 2). For the Oaxaca crosses, we found that the genomes of F2 progeny were 1.15 times larger than the B73 genome and that the BC1F2 progeny were 1.2 times larger. For the *Zea luxurians* crosses, the genomes of F2 progeny were 1.31 times larger than the B73 genome and the BC1F2 progeny were 1.47 times larger. The seven fully assembled B73 centromeres segregate in these progeny, providing the opportunity to measure changes in CENH3 area as a function of genome size in the context of identical centromere sequences.

**Figure 3.**
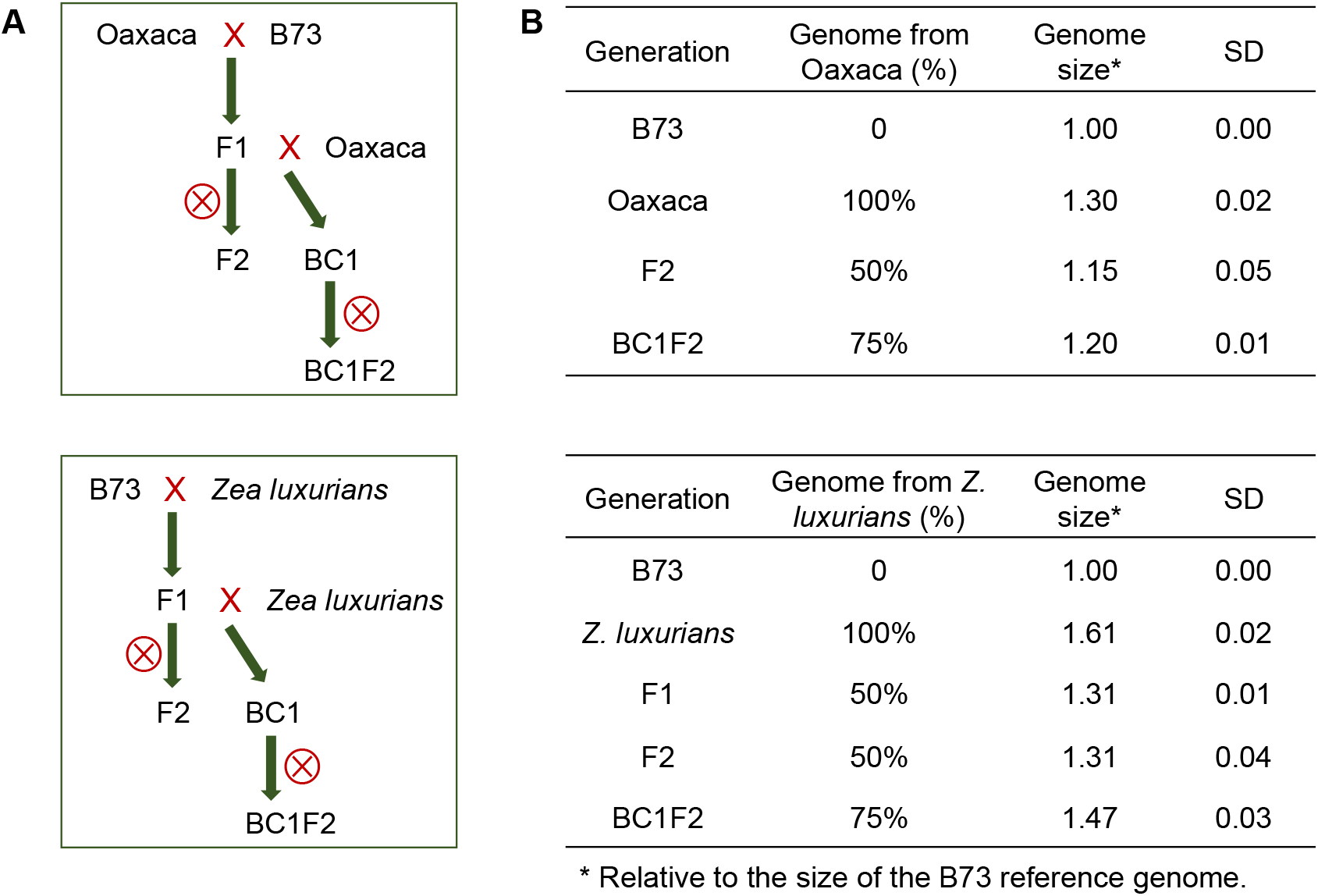
Crossing scheme for generating maize lines with different genome sizes. (A) Crossing schemes for transferring B73 centromeres into the Oaxaca and *Zea luxurians* backgrounds. (B) Genome sizes of B73, Oaxaca, *Zea luxurians* and their hybrids. Genome sizes are averages from 3-11 plants per family. SD indicates standard deviation.

#### B73 X Oaxaca hybrids

While the Oaxaca genome has not been sequenced, CENH3 ChIP-seq revealed that multiple centromeres from Oaxaca are similar to those in B73 (Supplemental Figure 2A). To avoid complexities associated with mapping two centromeres onto a single reference, we developed PCR markers to distinguish Oaxaca centromeres and focused our analysis entirely on B73 centromeres that were homozygous in the F2 or BC1F2 progeny (Supplemental Table 3). Analysis of the data revealed that four of the B73 centromeres examined (2, 4, 8 and 10) were significantly larger in the F2 and BC1F2 Oaxaca backgrounds than in their original smaller-genome context, and that the increases were correlated with genome size (Figure 4, Supplemental Table 4). The single exception was centromere 9 which had a statistically similar size in the B73 and the Oaxaca F2 progeny.

**Figure 4.**
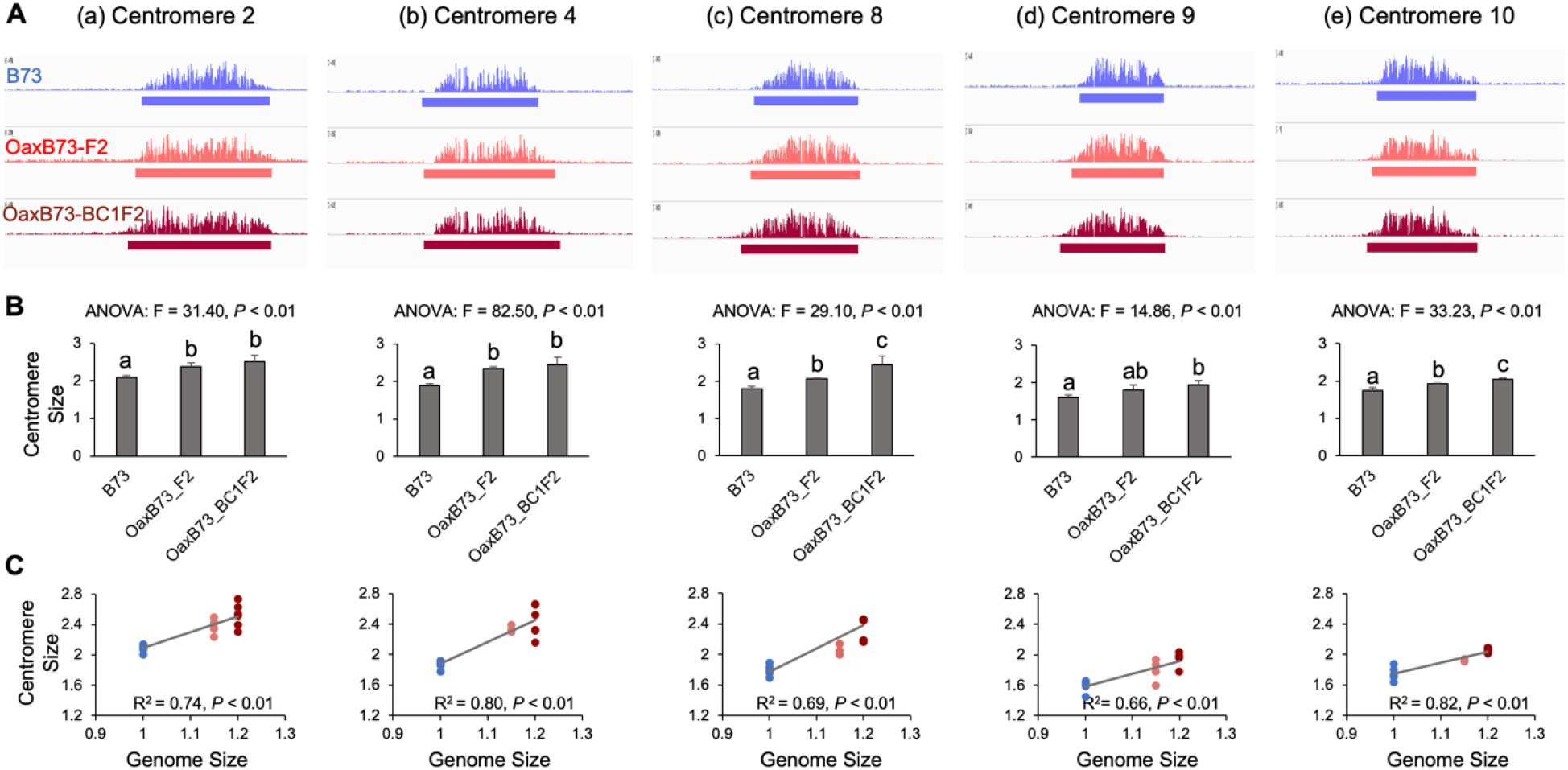
Centromeres are expanded in B73 X Oaxaca F2 and BC1F2 progeny. (A) CENH3-ChIP profiles of B73, Oaxaca X B73 F2 and Oaxaca X B73 BC1F2 progeny for five centromeres. The window size is 5Mb. (B) ANOVA analysis of centromere sizes across different lines. Bar graphs show mean centromere size comparison among different lines. Letters represent different groups that are statistically different (*P* < 0.05). Centromere sizes in the F2 and BC1F2 progeny were significantly larger than in B73, with the sole exception of centromere 9 in the F2 generation. Bars represent SD. (C) Linear regression analyses of centromere sizes across different lines. Dots represent different individuals. Blue: B73, orange/red: F2, dark red: BC1F2. Genome sizes are averages based on 3-11 individuals. The unit of centromere size is Mb.

#### *B73 X* Zea luxurians *hybrids*

The centromeres in *Zea luxurians* are known to contain long CentC arrays on every centromere (Albert *et al.* 2010). ChIP data from *Zea luxurians* consistently yielded high enrichment for CentC, but when these data were mapped to the B73 reference, there were no clear peaks (Supplemental Figure 2B). This is because uniquely-mapping reads generally exclude simple tandem repeats such as CentC. The absence of significant ChIP-seq read alignment from *Zea luxurians* centromeres allowed us to assess B73 centromere size in both the heterozygous and homozygous conditions.

The data reveal that in first generation F1 hybrids between B73 and *Zea luxurians*, there were little or no changes in centromeres size. We observed only minor increases in the sizes of Cen4, Cen5 and Cen8 in F1 progeny and no obvious changew in Cen2, Cen3, Cen9 and Cen10 (Figure 5). The differences were more pronounced in the F2 progeny where Cen2, Cen3, Cen4, Cen5 and Cen8 were significantly larger and Cen9 and Cen10 showed positive trends that were not statistically significant (Figure 5, Supplemental Table 5). An example is centromere 5, which is 1.86 Mb in the B73 inbred but expanded to 2.4 Mb in the B73 X *Zea luxurians* F2 progeny. Although there were only three B73 centromeres segregating in the BC1F2 population (Cen5, Cen9 and Cen10), all three were significantly expanded, confirming the trends observed in the F2 progeny. Taken together, the data from NAM centromere comparisons and Oaxaca and *Zea luxurians* crosses indicate that centromere size is positively correlated with genome size.

**Figure 5.**
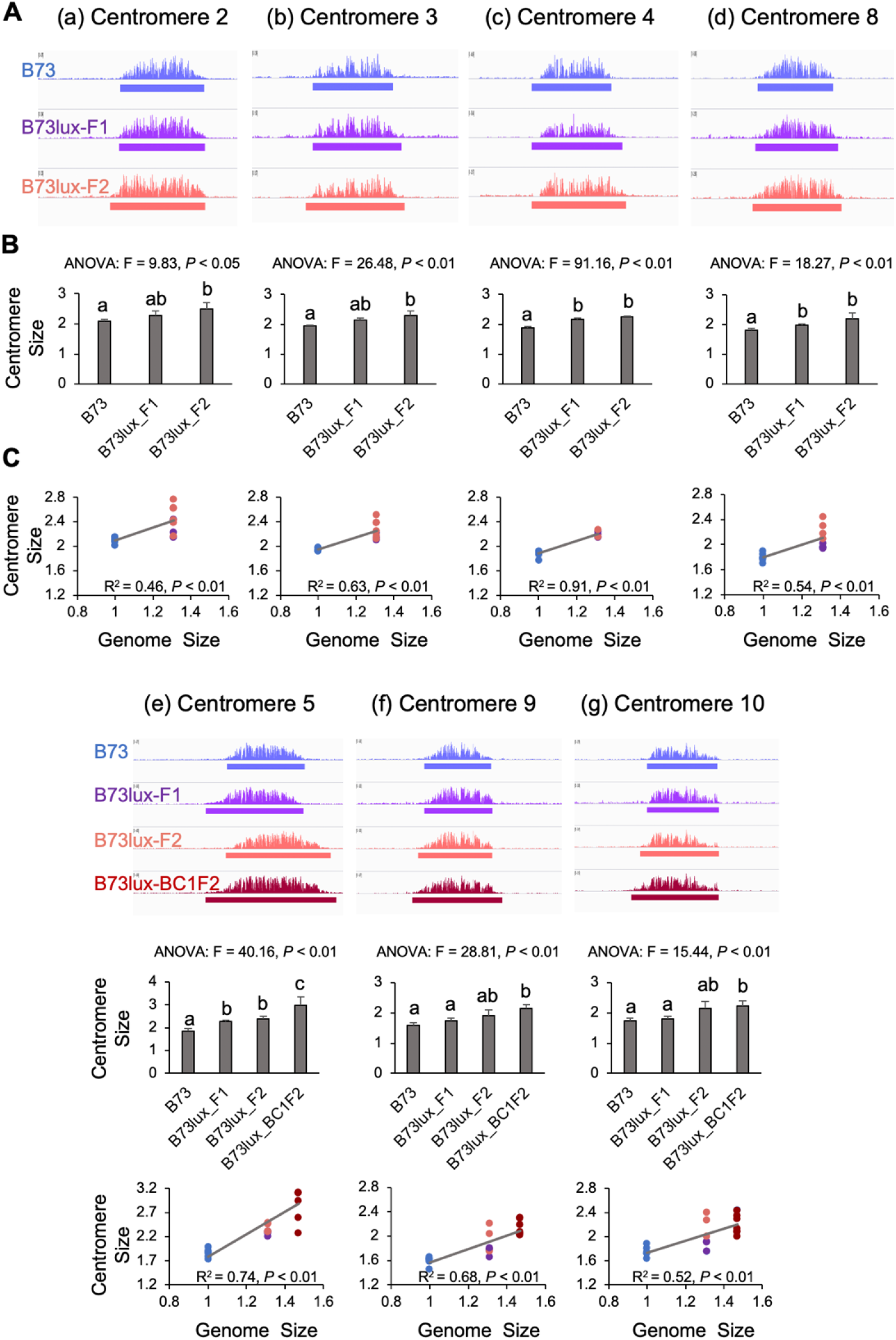
Centromeres are expanded in the B73 X *Z. luxurians* F2 and BC1F2 progeny. (A) CENH3-ChIP profiles of B73, B73 X *Z. luxurians* F1, F2 and BC1F2 progeny for seven centromeres. The window size is 5Mb. (B) ANOVA analysis of centromere sizes across different lines. Bar graphs show mean centromere size comparison among different lines. Letters represent different groups that are statistically different (*P* < 0.05). The sizes of Cen4, Cen5 and Cen8 in B73 X *Z. luxurians* F1 were significantly larger than B73, while the sizes of Cen2, Cen3, Cen9 and Cen10 in the F1 were not significantly larger. The sizes of Cen2, Cen3, Cen4, Cen5 and Cen8 in the F2 were significantly larger than B73, while the sizes of Cen9 and Cen10 in the F2 were not significantly larger than B73. The sizes of all centromeres in the BC1F2 were significantly larger than in B73. Bars represent SD. (C) Linear regression analyses of centromere sizes in F2 and BC1F2 progeny. Dots represent different individuals. Blue: B73, purple: F1, orange/red: F2, dark red: BC1F2. Genome sizes are averages based on 3-10 individuals. The unit of centromere size is Mb.

### Threefold overexpression of CENH3 does not affect centromere size

It is possible that centromere size is determined by the total amount of CENH3 that is available to bind to centromeric DNA. This hypothesis is supported by early work in Drosophila showing that overexpression of CENP-A/Cid caused a spreading of centromere locations to ectopic sites (Heun *et al.* 2006; Lacoste *et al.* 2014). A recent study of maize lines overexpressing a YFP-tagged version of CENH3 described subtle shifting of centromere locations, partially supporting this view (Feng *et al.* 2019). However, the YFP-tagged *CENH3* gene was not sufficient to complement a strong hypomorphic mutation, indicating it may not be fully functional or not expressed in all required cell types (Feng *et al.* 2019).

We were able to address this hypothesis using materials created for a study of how a *cenh3* null mutation behaves in crosses (Wang et al, *in press*). CRISPR-Cas9 was used to create a stop codon in the N-terminal tail of the *cenh3* gene. As a means to propagate the null, we also introduced a complete genomic copy of the transgene that differs from wild type by five silent nucleotide changes. RNA-seq of one of the transgenic lines (CENH3-Ox-1) showed approximately four-fold higher expression of the transgene than the wild type copy of *CENH3* (Figure 6B). Quantitative PCR analysis of genomic DNA from the CENH3-Ox-1 line suggested that the high *CENH3* expression in this line is caused by multiple transgene insertions (Figure 6B), which is a frequent occurrence in *Agrobacterium* transformants (Shou *et al.* 2004; Jupe *et al.* 2018). These four transgenes are sufficient to fully complement the *cenh3* null mutation (Wang et al, *in press*).

**Figure 6.**
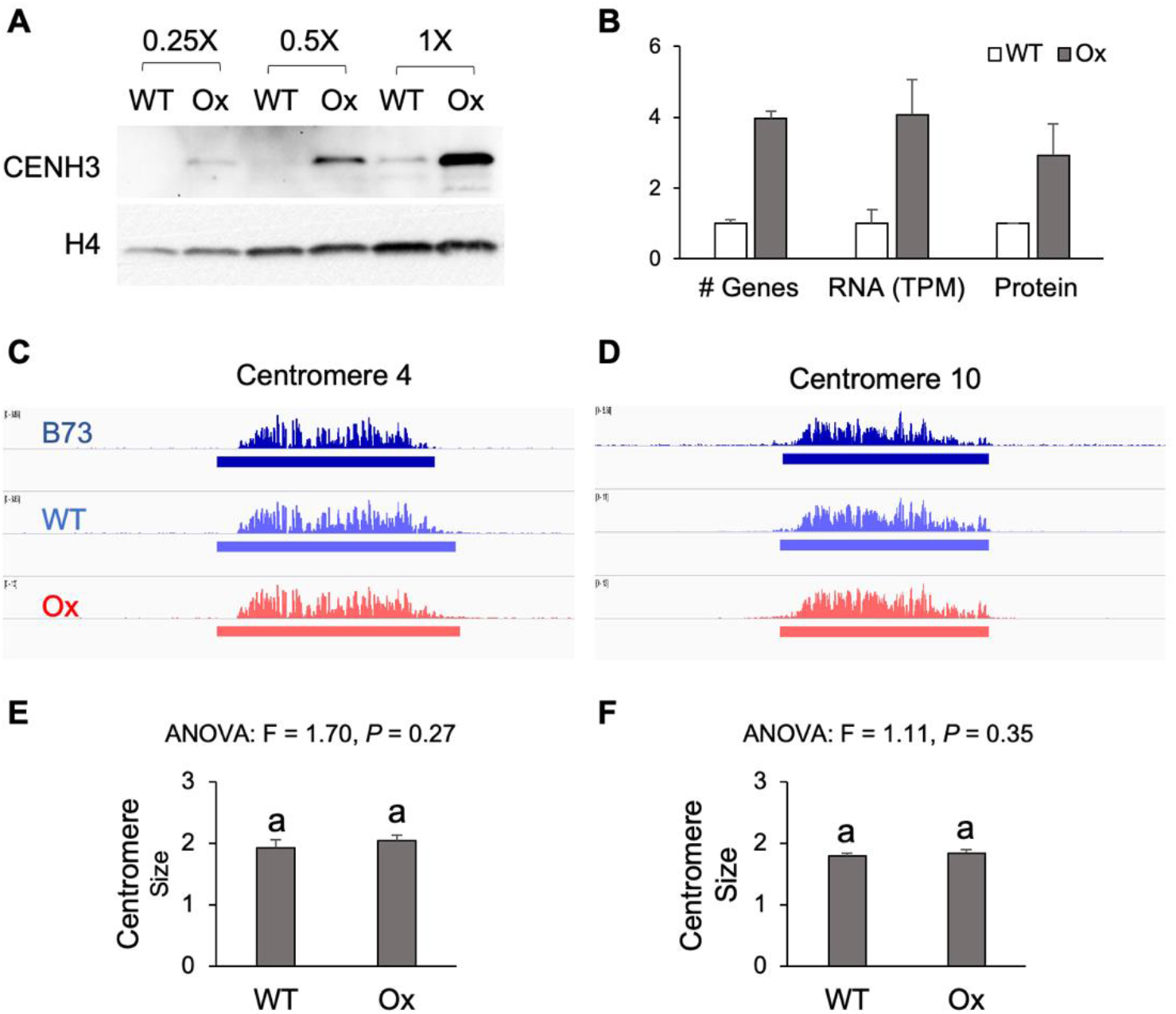
Centromere size is stable in the CENH3-overexpression lines. (A) Protein blot analysis of maize CENH3 expression levels in roots of wild type (WT) and CENH3-Ox-1 (Ox) lines. Nuclear protein was diluted to 0.25X, 0.5X and 1X. The same blot was incubated with antibodies to histone H4 as a loading control. (B) Quantification of *CENH3* gene copy number. DNA (gene) copy number was estimated by qPCR, mRNA expression was estimated from RNA-seq, and protein levels interpreted as the relative staining intensity of CENH3 and histone H4 on protein blots. Error bars show standard deviation. Wild type expression was set to one in each experiment. *CENH3* is a single copy gene in wild type lines. (C) and (D) CENH3 ChIP-seq profiles for B73, wild type siblings of the Ox lines, and Ox lines for centromeres 4 and 10. The window size is 5 Mb. (E) and (F) ANOVA analyses of centromere sizes across different lines. There were no significant differences in the sizes of Cen4 and Cen10 between WT and Ox (*P* < 0.05). The unit of centromere size is Mb. Bars in (B), (E) and (F) represent SD.

Analysis of leaf and root protein revealed that CENH3-Ox-1 lines have approximately three-fold higher nuclear CENH3 levels than wild type lines (Figure 6A, 6B), providing excellent material to test whether altered CENH3 levels change centromere size. The transgenic lines have a mixed genetic background but centromeres 4 and 10 are identical to those in B73. CENH3 ChIP-seq analysis of these two centromeres revealed no significant size differences between CENH3-Ox-1 lines and wild type siblings or the B73 inbred (Figure 6C-F). We also did not observe any new peaks in non-centromeric regions. These data indicate that centromere size is not affected by a threefold increase in CENH3 protein levels.

## Discussion

Here we combine data from a large collection of centromere sequences, empirical manipulations of genome size, and a novel CENH3 overexpression line to test how maize cells determine centromere size. Data from many sources suggest that at least in plants, DNA sequence alone does not determine centromere location. Here, viewed from a total centromere size perspective, we again find that centromere sequence does not itself determine the distribution of CENH3 (Figure 2B). Comparisons among the fully assembled centromeres in NAM inbreds revealed a weak positive correlation between total genome size and centromere size (Figure 2A) as predicted from earlier work (Zhang and Dawe 2012; Wang *et al.* 2014). The weakness of the correlation suggests that other factors also influence centromere size, possibly including small sequence motifs can have an effect on local positioning of CENH3 nucleosomes (Gent *et al.* 2011). In the oat-maize addition experiment, some centromeres expanded unidirectionally, suggesting barriers to CENH3 encoded in the DNA (Wang *et al.* 2014). It is also possible that our observed correlation between chromosome size and centromere size (Figure 2B) reflects inherent structural constraints encoded in the DNA.

CENP-A/CENH3 binds directly to DNA and is widely interpreted as a limiting factor for centromere establishment. Early work in Drosophila demonstrated that excess CENP-A was inappropriately distributed along chromosome arms, where it was sufficient to recruit all overlying kinetochore proteins and activate spurious centromeres (Heun *et al.* 2006). Similarly, overexpression of human CENP-A resulted in mislocalization of centromeric proteins and chromosome instability (Van Hooser *et al.* 2001; Shrestha *et al.* 2017). However, we found that in maize, overexpression of native CENH3 protein by threefold had no discernible effect on the size or distribution of CENH3 as assayed by ChIP (Figure 6). These results suggest that other factors limit the incorporation of excess CENH3 in maize, with likely candidates being the histone chaperones that direct CENH3 to centromeric locations (Mizuguchi *et al.* 2007; Dunleavy *et al.* 2009; Foltz *et al.* 2009; Chen *et al.* 2014). Among these are KNL2, which is required for CENH3 deposition in Arabidopsis (Lermontova *et al.* 2013) and NASP^SIM3^, which modulates soluble CENH3 levels (Le Goff *et al.* 2020). Another potential limiting factor is CENP-C, a key inner centromere protein that has been implicated in multiple aspects of centromere specification and stability (Du *et al.* 2010; Mitra *et al.* 2020).

Prior results demonstrated that when maize centromeres were transferred into the oat genome, their sizes increased roughly two-fold, in line with expectations based on the difference in genome size (Zhang and Dawe 2012; Wang *et al.* 2014). However, the wide oat-maize cross rarely succeeds and does not result in a stable hybrid (Kynast *et al.* 2001). Here we took a different approach of making natural crosses within *Zea* and tracking changes in centromere size over several generations. The results demonstrate that B73 centromeres increase in size when crossed into the larger Oaxaca and Z. *luxurians* backgrounds. Changes were clearly evident in F2 as well as BC1F2 progeny, revealing heritable increases (Figures 4 and 5). However, the size increases were less apparent in the first generation B73 X *Zea luxurians* individuals, consistent with prior results with the same F1 cross (Gent *et al.* 2017) and ruling out the alternative explanation that hybridization itself explains the increased centromere size. Multiple cellular generations or passage through meiosis may be required for CENH3 and associated binding partners and redistribute along chromosomes and reach a new centromere size equilibrium. Taken together, our results indicate that centromere size is scalable and responsive to changes in genome or cell size.

The impact of genome size on centromere size can be explained as a general cellular scaling process. Many forms of evidence from multiple species show strong correlations between genome size, nuclear size, and cell size (Price *et al.* 1973; Gregory 2001; Cavalier-Smith 2005; Gillooly *et al.* 2015; Robinson *et al.* 2018). With remarkably few exceptions, the entire cellular system scales in response to changes in genome size (Gregory 2001; Schmoller and Skotheim 2015; Amodeo and Skotheim 2016). These trends have been explained as an outcome of the fact that larger genomes are packaged into larger nuclei, and that nuclear size is at least indirectly correlated with cell size (Gregory 2001). Larger cells have more protein per cell and more and larger macromolecular structures such as mitochondria, microtubules, and ribosomes (Schmoller and Skotheim 2015). Given that the number of centromeres is constrained by the number of chromosomes, any increases in centromere size will be manifested as extensions of existing centromeres spread over larger chromosomal areas.

The scaling model not only requires scalable centromeres, but a deposition mechanism that is responsive to the amount of soluble precursors. A prior study of CENP-A dynamics in human cells provides support for the view that a mass-action mechanism regulates the number of CENP-A molecules bound to DNA (Bodor *et al.* 2014). The authors showed that about 4% of total CENP-A binds to centromeres over a range of natural expression variation, implying that centromere sizes vary with the amount of CENP-A available to bind (Bodor *et al.* 2014). They also overexpressed human CENP-A by approximately ~2.5 fold but did not observe corresponding changes in the amounts of the conserved kinetochore proteins CENP-C or NDC80, suggest that these and/or other key kinetochore proteins are limiting and help to buffer the effects of CENP-A overexpression. The available information from both human and maize show that while centromere sizes are malleable, moderate overexpression of CENP-A/CENH3 alone does not alter the size of the functional centromere domain, consistent with the view that multiple limiting factors together contribute to a stable centromere size equilibrium.

## Materials and Methods

### Plant materials and crossing

The plant materials used in this study were obtained from the Germplasm Resources Information Network (GRIN), Ames, Iowa. The lines were B73 (PI 550473), a domesticated landrace from Oaxaca, Mexico (PI 628470) and *Zea luxurians* (PI 462368). Crosses among lines were made over several years in the UGA Plant Biology greenhouses or an adjoining outdoor field site.

### NAM genome assemblies

Methodology for the PacBio assembly of NAM genomes is described in (Liu *et al.* 2020), with the exception that Nanopore data were not used. The descriptions and interpretations of these data are not yet published, but full assemblies and annotations are freely accessible at www.maizegdb.org/NAM_project.

### ChIP-seq

Whole seedlings including roots were collected from inbreds B73, CML103, CML277, CML333, HP301, IL14H, Ki11, Ki3, NC350, Oh7b, P39 and Tzi8 and CENH3 ChIP conducted as described previously (Gent *et al.* 2017) with the following modifications: During the nuclei extraction, we did not cut off pipet tips, and during micrococcal nuclease digestion, we used 2 μl micrococcal nuclease per 50 μl pelleted nuclei. For the overnight antibody incubation, we used μg of anti-maize CENH3 antibodies (Zhong et al 2003) and anti-rice CENH3 antibodies (Nagaki *et al.* 2004). During sequencing library preparation with the KAPA Hyper Prep kit (KK8502), we used a double-sided size selection with Mag-Bind® TotalPure NGS magnetic beads rather than a post-PCR gel purification. We removed large fragments with a 0.6X Mag-Bind bead cleanup (by adding 66 μL of beads to the 110 μL of ligation product and discarding the pellet), then removed small fragments with a 0.8X bead cleanup (by adding 136 μL of beads to the resulting 170 μL of supernatant from the last step and then discarding the new supernatant). All ChIP libraries were amplified with 5 cycles of PCR. Multiple adapters were used for pooling libraries (KAPA Single-Indexed Adapter Kit KK8700 and NEBNext^^®^^ Multiplex Oligos for Illumina NEB #E7535S/L). The DNA samples were sequenced using the Illumina NextSeq 500 platform and 150-nucleotide single-end reads were generated. The Sequence Read Archive run IDs for all the ChIP data of this study are listed in Supplemental Table 6.

PE100 Illumina CENH3 ChIP-seq reads for lines B97, CML228, CML322, CML247, CML52, CML69, KY21, MO18W, M37W, M162W, MS71, NC358, Oh43, and Tx303 were published previously (Schneider *et al.* 2016). The data were obtained from Genbank (SRP067358) and converted to single-end format using seqtk (https://github.com/lh3/seqtk).

### Measuring centromere size

The efficiency of CENH3 chromatin immunoprecipitation varies from day to day and sample to sample. The results are generally assessed after the experiment is over, when the average read depth within centromere cores is compared to the average read depth over chromosome arms. We observed previously (Gent *et al.* 2015), as well as in the current datasets, that measured centromere sizes vary with ChIP efficiency. This is because CENH3 ChIP-seq profiles take on the shape of bell-shaped curves. When there is low efficiency, a smaller profile of the curve exceeds the enrichment cutoff used to define edges of the centromere. To ameliorate this effect, we designed a custom workflow that amplifies ChIP-seq signals in the centromere relative to the chromosome arms, and consequently sharpens the centromere curve for samples with low efficiency (Supplemental Figure 1). The workflow involved four steps:

#### 1) Input data normalization

PE150 genomic input reads of all NAM lines (www.maizegdb.org/NAM_project) were subsampled to 30x with seqtk (v1.2, https://github.com/lh3/seqtk) relative to assembly size. The CENH3 ChIP-seq data in the form of PE100 reads were downloaded from SRP067358, converted to single-end data, and subsampled to 5 million reads using seqtk (v1.2). The SE150 ChIP reads generated in this study were subsampled to 3.33 million. Subsampled ChIP and genomic data were subjected to adapter removal with trimglore (v0.4.5, https://github.com/FelixKrueger/TrimGalore/) and mapped to corresponding genomes with bwa-mem (v0.7.17) at default parameters (Li and Durbin, 2009). PCR Duplicates were removed from bam files using piccard (v2.16) and alignments with a mapping quality higher than 20 were extracted with samtools (v1.9) (Li *et al.* 2009). The resulting CENH3 ChIP bam files were then normalized against input with deeptools (v3.2.1) (Ramírez *et al.* 2014) using the RPKM method with 5Kb non-overlapping windows (--binSize 5000 --normalizeUsing RPKM --scaleFactorsMethod None). Regions with an enrichment higher than 5 were extracted and merged into islands with bedtools (v2.28) (Quinlan 2014).

#### 2) Normalization of ChIP efficiency

Centromeres were located manually and placed into 5 Mb windows, then all remaining genomic space classified as background. The ratio between the sum of ChIP RPKM values (>=0) in the 5 Mb centromere regions (core) and background areas (all non-core areas) were used to calculate ChIP enrichment (Supplemental Fig1A). The core/background ratios were then modified using the formula (1+ core/background) × ChIP RPKM enrichment. This scaling step amplified signal in the centromere relative to chromosome arms. The resulting ChIP bedgraph files exhibited more pronounced curves compared with that before scaling (Supplemental Fig1C).

#### 3) Merging ChIP islands separated by alignment gaps

Highly repetitive regions result in either an absence of aligned reads or the alignment of far more reads than expected. Alignment gaps were defined as regions (>100bp) with lower than 2 or higher than 101 reads mapped using bedtools (v2.28). The resulting gaps resulted in isolated “ChIP islands” that we presumed would be connected if not for the intervening gap. Islands within 100 kb of each other that exceeded the 5-fold enrichment cutoff were merged using bedtools merge (v2.28; −d 15000). After this initial merging step, islands larger than 15 kb with enrichments higher than 3 were merged with a 50 kb interval using bedtools merge (v2.28; −d 50000).

#### 4) Using replicates to remove outliers

For lines with ChIP replicates, final coordinates were determined and centromere sizes were calculated for each replicate. Centromere sizes were compared among replicates, and outliers were removed using the mean absolute deviation (MAD) method. The mean centromere sizes among replicates were calculated after outlier removal.

Using these methods we compared centromere sizes across four different IL14H biological replicates. While the ChIP enrichment ranged from 7.36 to 13 (fold increase over background in the 5Mb core region), their measured centromere sizes after processing were similar (Supplemental Figure 1C).

### CRM and CentC annotation

CRM elements were identified by aligning complete copies of CRM1, CRM2, CRM3, and CRM4 (Sharma and Presting 2008) to the assemblies using BLAST (v2.2.26). The output was filtered by applying a 150 bp alignment length cutoff and enforcing an E-value lower than 0.0001. CentC was identified by aligning a consensus sequence (Gent *et al.* 2017) using BLAST and applying a 30 bp alignment length cutoff and enforcing an E-value lower than 0.001. The amount of CentC and CRM in active centromeres and chromosome arms were then quantified using bedtools merge and bedtools intersect from bedtools (v2.28).

### Genome size measurements of Oaxaca and *Zea luxurians* lines

Genome sizes were estimated by flow cytometry. Young leaf samples from single plants were sent to Plant Cytometry Services (Schijndel, the Netherlands) for flow cytometry measurements using *Vinca major* (2C = 4.2 pg) as an internal standard. We also included the reference maize inbred B73 in each batch as a second internal control to reduce technical error. Genome sizes were divided by the size of the B73 genome (where B73 was assigned a value of 1.0). Genome sizes measured in this study are listed in Supplemental Table 2.

### B73-Oaxaca, B73-luxurians hybrids and *CENH3* transgenic lines centromere analysis

CENH3 ChIP-seq SE150 data from the Oaxaca-B73, *Zea luxurians*-B73 and CENH3-Ox-1 overexpression lines described here were subsampled to 3.33 million reads. Additional ChIP and input samples from the parental Oaxaca and *Zea luxurians* lines were downloaded from SRP105290 (Gent et al., 2017). The subsampled reads were trimmed with TrimGalore (version 0.4.5) and mapped to Zm-B73-REFERENCE-NAM-5.0 (https://nam-genomes.org/) with BWA-mem (version 0.7.17) at default parameters (Li and Durbin, 2009). Only uniquely mapped reads (defined with MAPQ scores of at least 20) were used for peak calling. Centromere sizes were determined using the same methods used for NAM centromere analysis, except that B73 30X genomic Illumina data were used as input reads for all samples. After the merging steps, small islands less than 100Kb were manually removed. The results were visualized using IGVTools (version 2.3.98) at coverage calculated on 5kb intervals (Thorvaldsdóttir *et al.* 2013).

### Centromere genotyping

Mapping ChIP-seq reads from Oaxaca to B73 revealed that Oaxaca centromeres 2, 3, 8 and 9 have similar locations as the centromeres in B73 (Supplemental Figure 2). Using ChIP-seq reads, we identified SNPs with GATK HaplotypeCaller (v3.8-1) at default parameters (Poplin *et al.*). SNP2CAPS software (Thiel *et al.* 2004) was used to design Cleaved Amplified Polymorphic Sequence (CAPS) markers that distinguish B73 centromeres 2, 3, 8, 9 from those in the Oaxaca line (Supplemental Table 3). Young leaf DNA was prepared (Clarke 2009) and PCR products digested with restriction enzymes (Supplemental Table 3) to identify lines homozygous for the B73 centromeres.

In Oaxaca-B73 F2 and BC1F2 progeny, we genotyped the samples directly using ChIP-seq data. ChIP-seq reads were mapped to Zm-B73-REFERENCE-NAM-5.0 with BWA-mem (version 0.7.17) at default parameters. The results were then used for SNP calling with GATK HaplotypeCaller at default parameters. If no SNPs were present in the centromeric region, the corresponding centromere was classified as homozygous for the B73 centromere.

### Measuring *CENH3* copy number and expression in CENH3-Ox-1 lines

We prepared the CENH3-Ox-1 line by transforming maize Hi-II with the construct *gRNA-ImmuneCENH3* using *Agrobacterium* mediated transformation (Wang et al, *in press*). The *ImmuneCENH3* gene contains 6454 bp of the native CENH3 gene (coordinates Chr6:166705239-166711693 on Zm-B73-REFERENCE-NAM-5.0) with five silent codon changes to render it immune to a transacting guide RNA. The promoter includes 2184 bp of sequence upstream of the ATG. The transgene is sufficient to fully complement a *cenh3* null allele.

Quantitative PCR was used to determine *CenH3* gene copy number. Young leaf DNA was prepared (Clarke 2009) from three biological replicates from wild type and CENH3-Ox-1 transgenic lines. qPCR was carried out using a BioRad CFX96 Real-Time PCR system using a SYBR Green qPCR kit (Thermo Fisher Scientific). The single copy *Adh1* gene (Osterman and Dennis 1989) was used as an internal control gene. Primers are listed in Supplemental Table 3.

For RNA-seq, mRNA was prepared from young leaves of three wild type and three overexpression lines using a plant total RNA kit (IBI Scientific IB47342). 800 ng of total RNA was used for library construction with a mRNA-seq kit (KAPA mRNA hyper prep kit #KK8580). RNA-seq reads were trimmed with Trimmomatic at the following parameters: LEADING:3 TRAILING:3 SLIDINGWINDOW:4:15 MINLEN:36 (Bolger *et al.* 2014), then the trimmed reads were mapped to Zm-B73-REFERENCE-NAM-5.0 with hisat2 at the following parameters: --min-intronlen 20, --max-intronlen 500000, --rna-strandness R (Kim *et al.* 2019). The alignments were converted to BAM files and sorted with SAMtools. Stringtie was used to compute gene expression levels using Transcripts Per Kilobase Million (TPM) = 1 as the cutoff (Pertea *et al.* 2015; Kim *et al.* 2019).

### Nuclear protein isolation and protein blotting

Approximately 2 g of flash-frozen leaves or roots were collected and chopped into 1.5 ml pre-chilled nuclei extraction buffer (1mM EDTA, 1x cOmplete™ Mini EDTA-free Protease Inhibitor Cocktail, 10 mM Tris-HCl pH 7.5, 10 mM NaCl, 0.2% NP-40, 5 mM 2-mercaptoethanol, 0.1 mM PMSF). The mixture was poured through miracloth and filtered through a 40 μm cell strainer. Then, 30 μl of the filtered sample was stained with 4,6-diamidino-2-phenylindole and nuclei counted using fluorescence microscopy. Nuclei concentrations were normalized based on these measurements. The nuclei were centrifuged at 5000 g for 5 mins and the pellets flash-frozen and stored at −80℃ until used for protein blots. Nuclei were resuspended in Laemmli buffer and loaded into 4-20% Mini-PROTEAN® TGX™ Precast Protein Gels (Bio-Rad Cat #4561093). SDS-PAGE and protein blotting were performed according to (Dawe *et al.* 2018). CENH3 was detected with anti-CENH3 antibodies (Zhong *et al.* 2002) (1:1000 dilution) and normalized to total H4 histones revealed by an anti-H4 antibodies (1:1000 dilution, Abcam, ab7311). Primary antibodies were detected using anti-rabbit secondary antibodies (1:5000 dilution, Anti-Rabbit IgG HRP Linked Whole Ab Sigma Cat# GENA934-1ML). The band intensities were quantified with Image J (Schneider *et al.* 2012).

## Statistical analysis

Welch’s analysis of variance (ANOVA) tests were performed to determine whether genome size and centromere size were significantly different across a variety of subgroups (Welch 1951). Pairwise comparisons among different subgroups were conducted with the available R package for Welch’s test (http://www.r-project.org; (Dag *et al.* 2018). Significance was set at P < 0.05. Linear regression analysis was performed with lm function from R to model the relationship between centromere size and genome size in individual groups (Rosner 2015).

## Data Availability

ChIP reads can be obtained from the NCBI Sequence Read Archive (ncbi.nlm.nih.gov/sra) under project PRJNA639705. Strains and reagents are available upon request.

## Acknowledgements

This work was funded by grant 1444514 from the National Science Foundation. We appreciate the support of the Georgia Genomics and Bioinformatics Core facility, **t**he Georgia Advanced Computing Resource Center, and the UGA Plant Biology greenhouse staff.

## Supplemental Figures

**Supplemental Figure 1.**
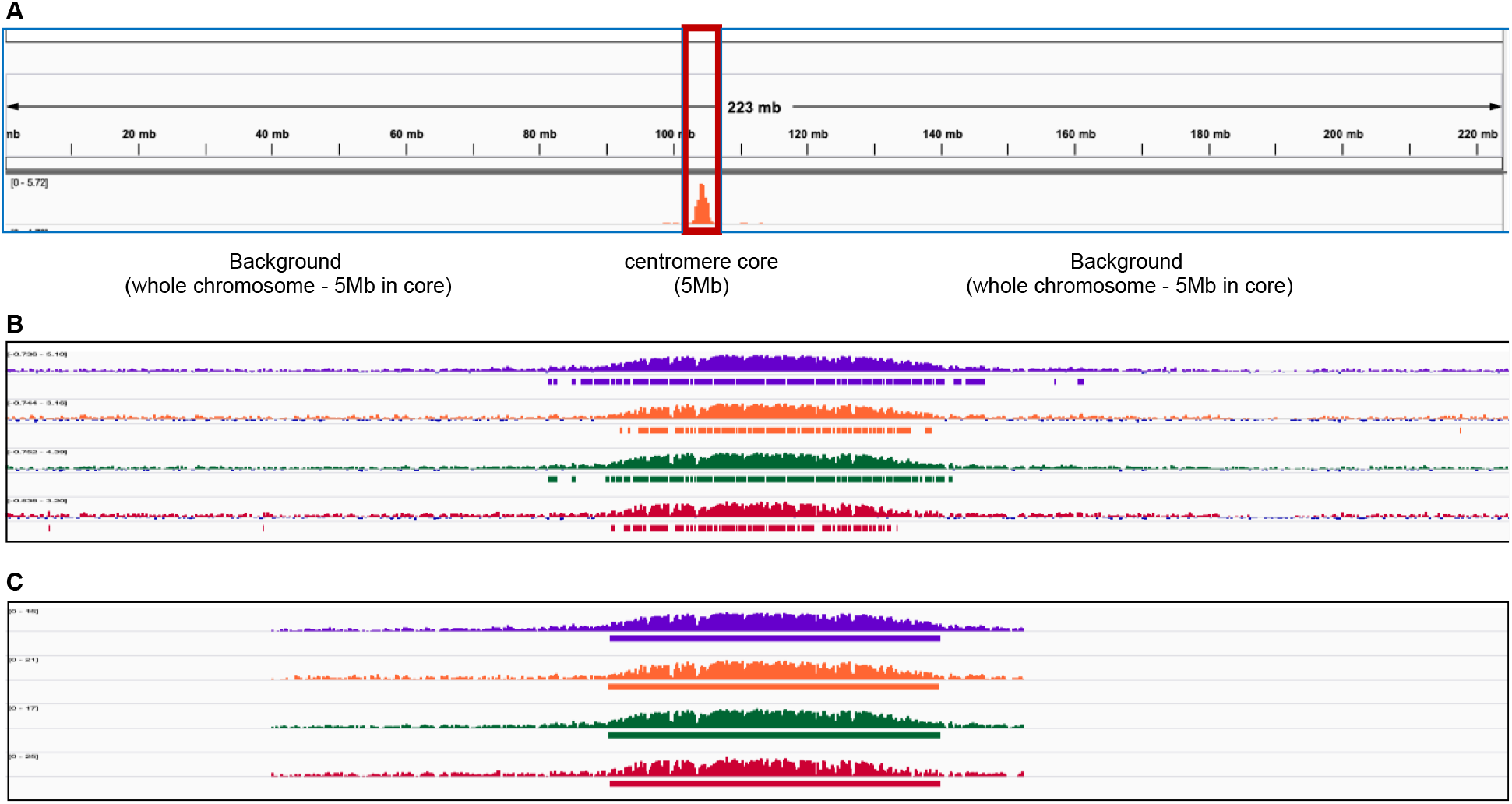
Impact of ChIP normalization on centromere size measurements. (A) The 5 Mb centromere core region for centromere 5 in the inbred IL14H. The regions around this 5 Mb domain were defined as background. (B) CENH3-ChIP profiles of four IL14H biological replicates of centromere 5 before scaling. (C) The same four biological replicates after normalizing for ChIP efficiency. The window size is 10 Mb.

**Supplemental Figure 2.**
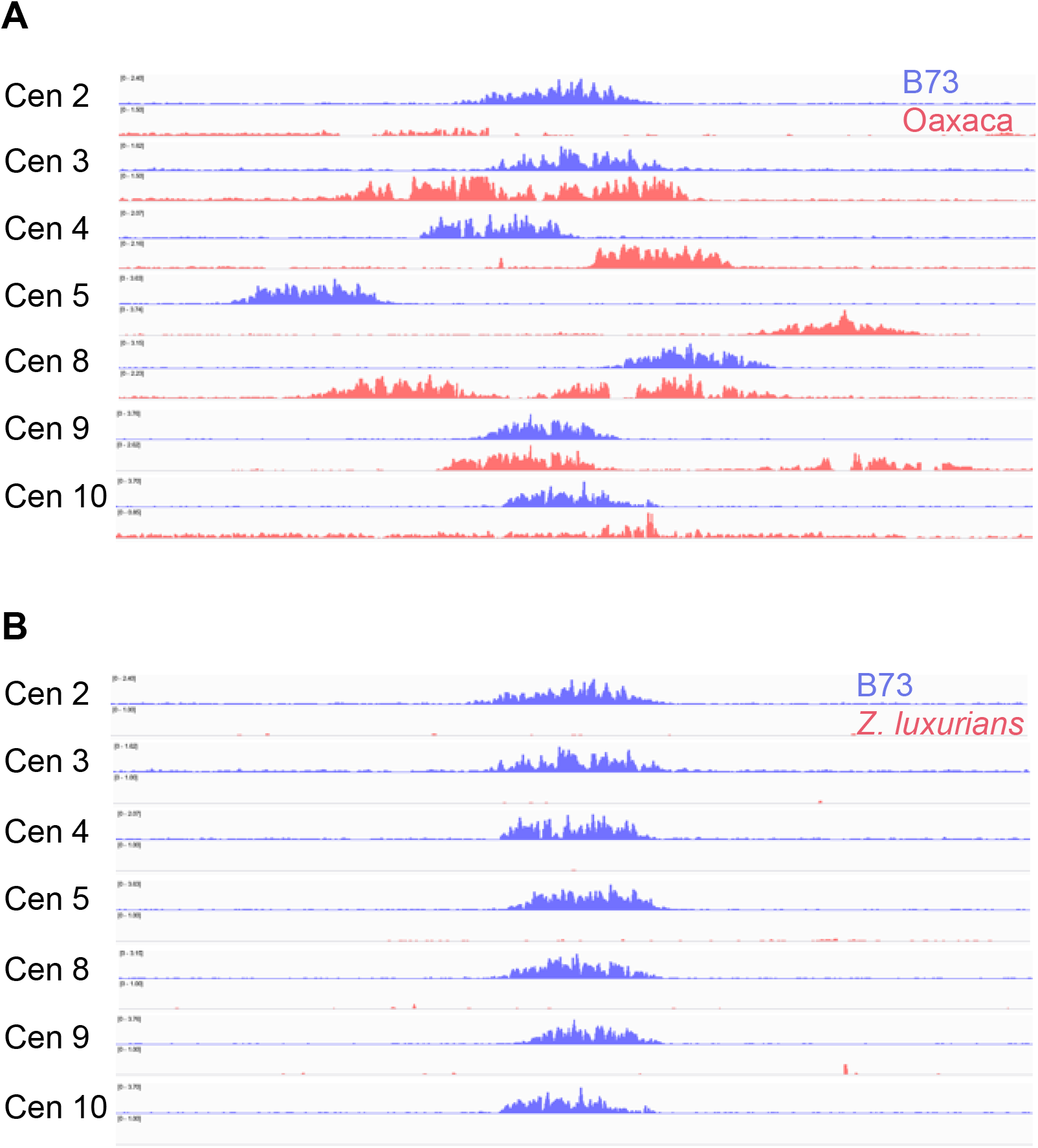
Comparison of B73 centromeres to those in Oaxaca and *Zea luxurians*. (A) CENH3 ChIP-seq profiles for seven B73 and Oaxaca centromeres mapped to the B73 reference. The data are from a single heterozygous plant. (B) CENH3 ChIP-seq profiles for seven B73 and *Zea luxurians* centromeres mapped to the B73 reference. The data are from a single heterozygous plant. There appears to be no centromeres in *Zea luxurians* because the centromeres are embedded in long CentC arrays that lack homology to the B73 reference. The window size is 10Mb.

**Supplemental Table 1.** ChIP datasets generated in this study.

| Sample_name | Species/subspecies | Cultivar | Tissue | Genome size (GB) | Genome size (ratio to B73) | # of replicate |
| --- | --- | --- | --- | --- | --- | --- |
| B73 | Zea mays mays | B73 | seedlings | 2.30 | 1 | 1 |
| CML103 <sup>1</sup> | Zea mays mays | CML103 | seedlings | 2.31 | 1.00 | 2 |
| CML277 <sup>1</sup> | Zea mays mays | CML277 | seedlings | 2.34 | 1.02 | 2 |
| CML333 <sup>1</sup> | Zea mays mays | CML333 | seedlings | 2.40 | 1.04 | 3 |
| HP301 <sup>1</sup> | Zea mays mays | HP301 | seedlings | 2.21 | 0.96 | 3 |
| IL14H <sup>1</sup> | Zea mays mays | IL14H | seedlings | 2.10 | 0.91 | 3 |
| Ki11 <sup>1</sup> | Zea mays mays | Ki11 | seedlings | 2.38 | 1.03 | 3 |
| Ki3 <sup>1</sup> | Zea mays mays | Ki3 | seedlings | 2.35 | 1.02 | 3 |
| NC350 <sup>1</sup> | Zea mays mays | NC350 | seedlings | 2.40 | 1.04 | 3 |
| Oh7b <sup>1</sup> | Zea mays mays | Oh7b | seedlings | 2.20 | 0.96 | 2 |
| P39 <sup>1</sup> | Zea mays mays | P39 | seedlings | 2.10 | 0.91 | 3 |
| Tzi8 <sup>1</sup> | Zea mays mays | Tzi8 | seedlings | 2.35 | 1.02 | 3 |
| OaxB73_F2 | Zea mays mays | B73 X PI 628470 | seedlings | 2.65 | 1.15 | 9 |
| OaxB73_BC1F2 | Zea mays mays | B73 X PI 628470 | seedlings | 2.76 | 1.20 | 15 |
| B73lux_F1 | Zea mays mays x Zea luxurians | B73 x PI 422162 | seedlings | 3.01 | 1.31 | 3 |
| B73lux_F2 | Zea mays mays x Zea luxurians | B73 x PI 422162 | seedlings | 3.01 | 1.31 | 6 |
| B73lux_BC1F2 | Zea mays mays x Zea luxurians | B73 x PI 422162 | seedlings | 3.38 | 1.47 | 9 |
| transgenic_WT | Zea mays mays | mixed | seedlings | NA | NA | 3 |
| transgenic_OX | Zea mays mays | mixed | seedlings | NA | NA | 3 |

<sup>1</sup> Genome sizes are from (Chia *et al.* 2012).

**Supplemental Table 2.**
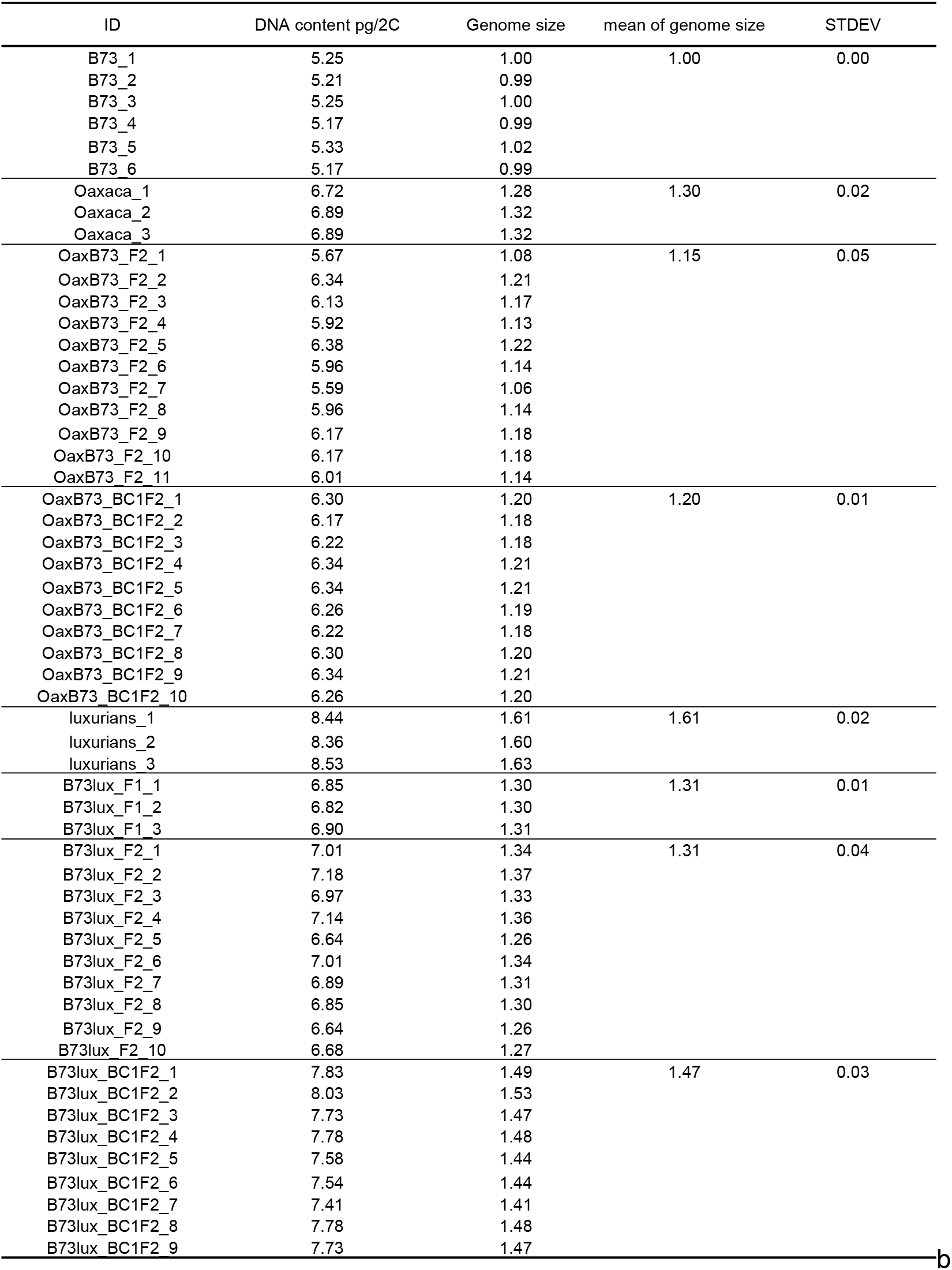
Genome sizes measured in this study. Measured genome sizes were divided by the average size of the six B73 samples.

**Supplemental Table 3.**
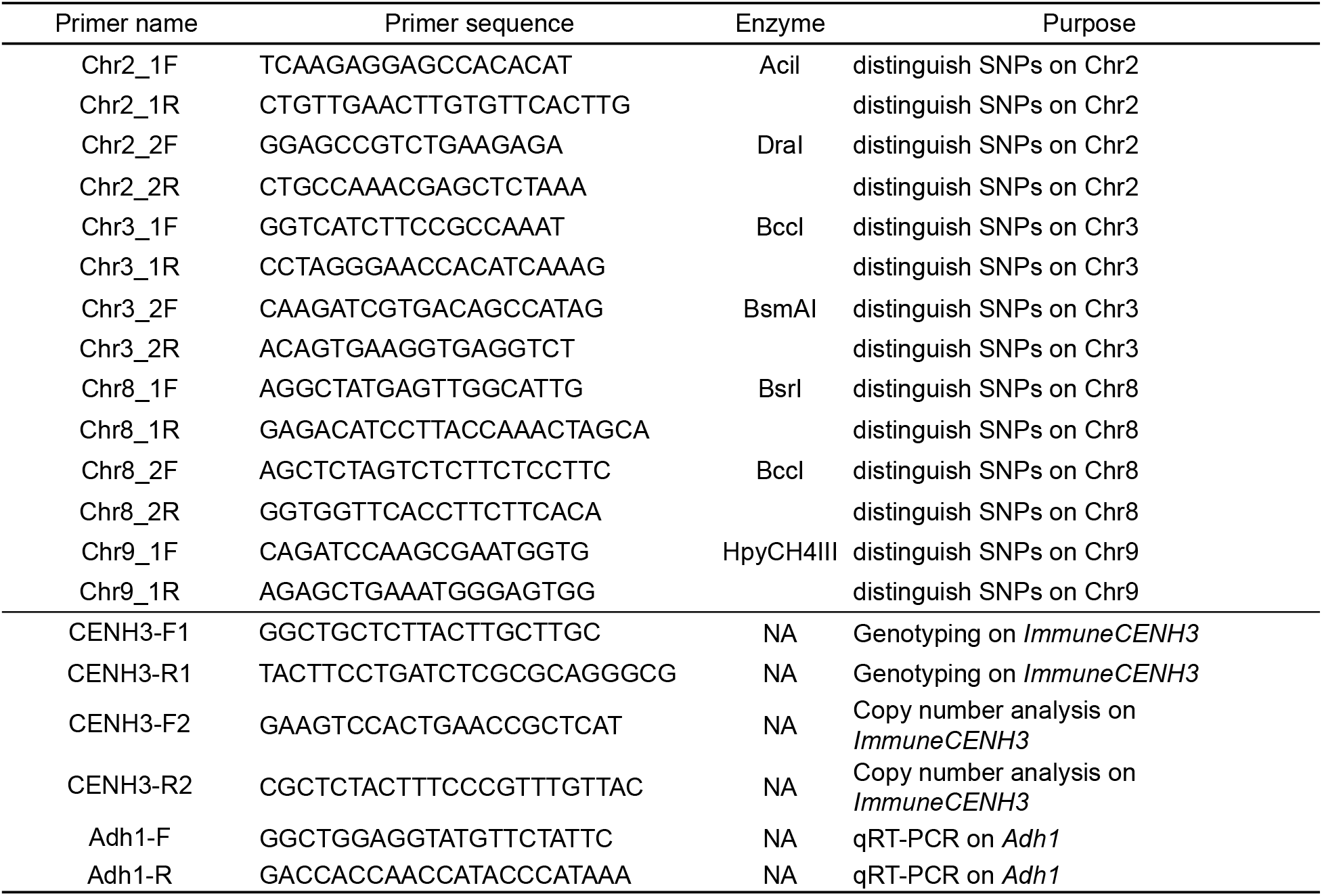
Primers used in this study.

**Supplemental Table 4.** Genome and centromere sizes in Oaxaca X B73 hybrids. Genome sizes were not measured for the plants subjected to ChIP. The genome sizes shown are averages of multiple sibling individuals measured from the same ears (Supplemental Table 2). For centromere sizes, the unit of measure is bp.

| ID | genome size | centromere2 | centromere3 | centromere4 | centromere8 | centromere9 | centromere10 |
| --- | --- | --- | --- | --- | --- | --- | --- |
| OaxB73_F2_1 | 1.15 | 2345000 | NA | NA | NA | NA | NA |
| OaxB73_F2_2 | 1.15 | 2420000 | 2219160 | NA | 1998403 | NA | NA |
| OaxB73_F2_3 | 1.15 | NA | NA | 2320000 | NA | 1870000 | NA |
| OaxB73_F2_4 | 1.15 | 2415000 | NA | NA | NA | NA | 1939465 |
| OaxB73_F2_5 | 1.15 | NA | NA | NA | NA | 1936000 | NA |
| OaxB73_F2_6 | 1.15 | 2491357 | NA | NA | NA | NA | NA |
| OaxB73_F2_7 | 1.15 | NA | NA | 2299125 | NA | NA | NA |
| OaxB73_F2_8 | 1.15 | NA | NA | NA | 2047653 | 1776712 | NA |
| OaxB73_F2_9 | 1.15 | 2240000 | NA | 2393658 | 2140354 | 1600000 | 1910817 |
| OaxB73_BC1F2_1 | 1.20 | 2732740 | 2578497 | 2159125 | NA | NA | 2015000 |
| OaxB73_BC1F2_2 | 1.20 | 2299486 | NA | NA | 2456832 | NA | NA |
| OaxB73_BC1F2_4 | 1.20 | NA | NA | NA | NA | NA | NA |
| OaxB73_BC1F2_6 | 1.20 | NA | 2395000 | NA | 2460000 | NA | NA |
| OaxB73_BC1F2_8 | 1.20 | NA | 2500000 | 2664999 | NA | NA | NA |
| OaxB73_BC1F2_9 | 1.20 | 2625000 | 2379160 | 2520000 | NA | NA | NA |
| OaxB73_BC1F2_10 | 1.20 | NA | NA | NA | NA | NA | NA |
| OaxB73_BC1F2_11 | 1.20 | NA | 2310000 | 2315000 | 2186832 | 1961778 | NA |
| OaxB73_BC1F2_12 | 1.20 | NA | NA | 2650000 | 2160821 | 1781712 | 2088648 |
| OaxB73_BC1F2_13 | 1.20 | NA | NA | NA | 2456832 | NA | NA |
| OaxB73_BC1F2_15 | 1.20 | NA | 2225287 | NA | 2441068 | NA | 2073648 |
| OaxB73_BC1F2_16 | 1.20 | NA | 2375000 | NA | 2900780 | 1986778 | NA |
| OaxB73_BC1F2_17 | 1.20 | 2520000 | NA | 2320000 | NA | 2036778 | 2038648 |
| OaxB73_BC1F2_18 | 1.20 | 2391577 | NA | NA | NA | NA | NA |
| OaxB73_BC1F2_19 | 1.20 | 2538991 | NA | NA | NA | NA | NA |

**Supplemental Table 5.** Genome and centromere sizes in B73 X *Zea luxurians* hybrids. Genome sizes were not measured for the plants subjected to ChIP. The genome sizes shown are averages of multiple sibling individuals measured from the same ears (Supplemental Table 2). For centromere sizes, the unit of measure is bp.

| ID | genome size | centromere 2 | centromere3 | centromere4 | centromere5 | centromere8 | centromere9 | centromere10 |
| --- | --- | --- | --- | --- | --- | --- | --- | --- |
| B73lux_F1_1 | 1.31 | 2445000 | 2100638 | 2200000 | 2265464 | 1965000 | 1765000 | 1752649 |
| B73lux_F1_2 | 1.31 | 2230000 | 2140287 | 2148658 | 2200505 | 1955000 | 1655000 | 1752649 |
| B73lux_F1_3 | 1.31 | 2150000 | 2214447 | 2154125 | 2315000 | 2020000 | 1795000 | 1905817 |
| B73lux_F2_1 | 1.31 | 2765000 | 2114450 | NA | 2303903 | 2445000 | 1776712 | 2275000 |
| B73lux_F2_2 | 1.31 | 2160482 | 2385000 | NA | 2457934 | 2175000 | 1760000 | NA |
| B73lux_F2_3 | 1.31 | 2426577 | 2520000 | NA | 2488026 | 1936832 | 1800000 | 1920817 |
| B73lux_F2_4 | 1.31 | 2620000 | 2180287 | 2240992 | NA | 2100000 | 2036778 | NA |
| B73lux_F2_5 | 1.31 | 2633991 | 2245000 | NA | NA | 2302376 | 2200000 | 2407741 |
| B73lux_F2_6 | 1.31 | 2381577 | 2379160 | 2269125 | 2313940 | NA | NA | 2008246 |
| B73lux_BC1F2_1 | 1.47 | NA | NA | NA | 3280000 | NA | 2296778 | NA |
| B73lux_BC1F2_2 | 1.47 | NA | NA | NA | 3125000 | NA | 2301778 | 2347913 |
| B73lux_BC1F2_6 | 1.47 | NA | NA | NA | NA | NA | 2055000 | 2435700 |
| B73lux_BC1F2_11 | 1.47 | NA | NA | NA | 3105000 | NA | 2021778 | 2295000 |
| B73lux_BC1F2_13 | 1.47 | NA | NA | NA | 2275000 | NA | NA | 2078648 |
| B73lux_BC1F2_15 | 1.47 | NA | NA | NA | 2937934 | NA | 2186778 | 2132913 |
| B73lux_BC1F2_16 | 1.47 | NA | NA | NA | 2597302 | NA | NA | 2005000 |
| B73lux_BC1F2_17 | 1.47 | NA | NA | NA | 3309471 | NA | 2035000 | 2435000 |
| B73lux_BC1F2_19 | 1.47 | NA | NA | NA | 3220000 | NA | 2065000 | 2095000 |

**Supplemental Table 6.** Sequence Read Archive run IDs for all ChIP data used in this study.

| Sample_name | Species/<br>subspecies | Cultivar | Accession | ChIP SRA run | input SRA<br>run | Library<br>layout | Tissue | Reference |
| --- | --- | --- | --- | --- | --- | --- | --- | --- |
| CML103_1 | Zea mays | CML103 | PRJNA639705 | SRR12023802 | NA | paired | seedling | this study |
| CML103_2 | Zea mays | CML103 | PRJNA639705 | SRR12023791 | NA | paired | seedling | this study |
| CML277_1 | Zea mays | CML277 | PRJNA639705 | SRR12023780 | NA | paired | seedling | this study |
| CML277_2 | Zea mays | CML277 | PRJNA639705 | SRR12023769 | NA | paired | seedling | this study |
| CML333_1 | Zea mays | CML333 | PRJNA639705 | SRR12023758 | NA | single | seedling | this study |
| CML333_2 | Zea mays | CML333 | PRJNA639705 | SRR12023747 | NA | single | seedling | this study |
| CML333_3 | Zea mays | CML333 | PRJNA639705 | SRR12023736 | NA | single | seedling | this study |
| HP301_1 | Zea mays | HP301 | PRJNA639705 | SRR12023730 | NA | paired | seedling | this study |
| HP301_2 | Zea mays | HP301 | PRJNA639705 | SRR12023729 | NA | paired | seedling | this study |
| IL14H_1 | Zea mays | IL14H | PRJNA639705 | SRR12023801 | NA | paired | seedling | this study |
| IL14H_2 | Zea mays | IL14H | PRJNA639705 | SRR12023800 | NA | paired | seedling | this study |
| IL14H_3 | Zea mays | IL14H | PRJNA639705 | SRR12023799 | NA | paired | seedling | this study |
| Ki11_1 | Zea mays | Ki11 | PRJNA639705 | SRR12023798 | NA | single | seedling | this study |
| Ki11_2 | Zea mays | Ki11 | PRJNA639705 | SRR12023797 | NA | single | seedling | this study |
| Ki11_3 | Zea mays | Ki11 | PRJNA639705 | SRR12023796 | NA | single | seedling | this study |
| Ki3_1 | Zea mays | Ki3 | PRJNA639705 | SRR12023795 | NA | paired | seedling | this study |
| Ki3_2 | Zea mays | Ki3 | PRJNA639705 | SRR12023794 | NA | paired | seedling | this study |
| NC350_1 | Zea mays | NC350 | PRJNA639705 | SRR12023793 | NA | paired | seedling | this study |
| NC350_2 | Zea mays | NC350 | PRJNA639705 | SRR12023792 | NA | paired | seedling | this study |
| Oh7b_1 | Zea mays | Oh7b | PRJNA639705 | SRR12023790 | NA | paired | seedling | this study |
| Oh7b_2 | Zea mays | Oh7b | PRJNA639705 | SRR12023789 | NA | paired | seedling | this study |
| P39_1 | Zea mays | P39 | PRJNA639705 | SRR12023788 | NA | single | seedling | this study |
| P39_2 | Zea mays | P39 | PRJNA639705 | SRR12023787 | NA | single | seedling | this study |
| P39_3 | Zea mays | P39 | PRJNA639705 | SRR12023786 | NA | single | seedling | this study |
| Tzi8_1 | Zea mays | Tzi8 | PRJNA639705 | SRR12023785 | NA | paired | seedling | this study |
| Tzi8_2 | Zea mays | Tzi8 | PRJNA639705 | SRR12023784 | NA | paired | seedling | this study |
| B73_4 | Zea mays | B73 | PRJNA639705 | SRR12023803 | NA | single | seedling | this study |
| OaxB73_F2_1 | Zea mays | OaxB73_F2 | PRJNA639705 | SRR12023783 | NA | single | seedling | this study |
| OaxB73_F2_2 | Zea mays | OaxB73_F2 | PRJNA639705 | SRR12023782 | NA | single | seedling | this study |
| OaxB73_F2_3 | Zea mays | OaxB73_F2 | PRJNA639705 | SRR12023781 | NA | single | seedling | this study |
| OaxB73_F2_4 | Zea mays | OaxB73_F2 | PRJNA639705 | SRR12023779 | NA | single | seedling | this study |
| OaxB73_F2_5 | Zea mays | OaxB73_F2 | PRJNA639705 | SRR12023778 | NA | single | seedling | this study |
| OaxB73_F2_6 | Zea mays | OaxB73_F2 | PRJNA639705 | SRR12023777 | NA | single | seedling | this study |
| OaxB73_F2_7 | Zea mays | OaxB73_F2 | PRJNA639705 | SRR12023776 | NA | single | seedling | this study |
| OaxB73_F2_8 | Zea mays | OaxB73_F2 | PRJNA639705 | SRR12023775 | NA | single | seedling | this study |
| OaxB73_F2_9 | Zea mays | OaxB73_F2 | PRJNA639705 | SRR12023774 | NA | single | seedling | this study |
| OaxB73_BC1F2_1 | Zea mays | OaxB73_BC1F2 | PRJNA639705 | SRR12023773 | NA | single | seedling | this study |
| OaxB73_BC1F2_2 | Zea mays | OaxB73_BC1F2 | PRJNA639705 | SRR12023772 | NA | single | seedling | this study |
| OaxB73_BC1F2_4 | Zea mays | OaxB73_BC1F2 | PRJNA639705 | SRR12023771 | NA | single | seedling | this study |
| OaxB73_BC1F2_6 | Zea mays | OaxB73_BC1F2 | PRJNA639705 | SRR12023770 | NA | single | seedling | this study |
| OaxB73_BC1F2_8 | Zea mays | OaxB73_BC1F2 | PRJNA639705 | SRR12023768 | NA | single | seedling | this study |
| OaxB73_BC1F2_9 | Zea mays | OaxB73_BC1F2 | PRJNA639705 | SRR12023767 | NA | single | seedling | this study |
| OaxB73_BC1F2_10 | Zea mays | OaxB73_BC1F2 | PRJNA639705 | SRR12023766 | NA | single | seedling | this study |
| OaxB73_BC1F2_11 | Zea mays | OaxB73_BC1F2 | PRJNA639705 | SRR12023765 | NA | single | seedling | this study |
| OaxB73_BC1F2_12 | Zea mays | OaxB73_BC1F2 | PRJNA639705 | SRR12023764 | NA | single | seedling | this study |
| OaxB73_BC1F2_13 | Zea mays | OaxB73_BC1F2 | PRJNA639705 | SRR12023763 | NA | single | seedling | this study |
| OaxB73_BC1F2_15 | Zea mays | OaxB73_BC1F2 | PRJNA639705 | SRR12023762 | NA | single | seedling | this study |
| OaxB73_BC1F2_16 | Zea mays | OaxB73_BC1F2 | PRJNA639705 | SRR12023761 | NA | single | seedling | this study |
| OaxB73_BC1F2_17 | Zea mays | OaxB73_BC1F2 | PRJNA639705 | SRR12023760 | NA | single | seedling | this study |
| OaxB73_BC1F2_18 | Zea mays | OaxB73_BC1F2 | PRJNA639705 | SRR12023759 | NA | single | seedling | this study |
| OaxB73_BC1F2_19 | Zea mays | OaxB73_BC1F2 | PRJNA639705 | SRR12023757 | NA | single | seedling | this study |
| B73lux_F1_1 | Zea mays subsp.<br>mays x Zea luxurians | B73lux_F1 | PRJNA639705 | SRR12023756 | NA | single | seedling | this study |
| B73lux_F1_2 | Zea mays subsp.<br>mays x Zea luxurians | B73lux_F1 | PRJNA639705 | SRR12023755 | NA | single | seedling | this study |
| B73lux_F1_3 | Zea mays subsp.<br>mays x Zea luxurians | B73lux_F1 | PRJNA639705 | SRR12023754 | NA | single | seedling | this study |
| B73lux_F2_1 | Zea mays subsp.<br>mays x Zea luxurians | B73lux_F2 | PRJNA639705 | SRR12023753 | NA | single | seedling | this study |
| B73lux_F2_2 | Zea mays subsp.<br>mays x Zea luxurians | B73lux_F2 | PRJNA639705 | SRR12023752 | NA | single | seedling | this study |
| B73lux_F2_3 | Zea mays subsp.<br>mays x Zea luxurians | B73lux_F2 | PRJNA639705 | SRR12023751 | NA | single | seedling | this study |
| B73lux_F2_4 | Zea mays subsp.<br>mays x Zea luxurians | B73lux_F2 | PRJNA639705 | SRR12023750 | NA | single | seedling | this study |
| B73lux_F2_5 | Zea mays subsp.<br>mays x Zea luxurians | B73lux_F2 | PRJNA639705 | SRR12023749 | NA | single | seedling | this study |
| B73lux_F2_6 | Zea mays subsp.<br>mays x Zea luxurians | B73lux_F2 | PRJNA639705 | SRR12023748 | NA | single | seedling | this study |
| B73lux_BC1F2_1 | Zea mays subsp.<br>mays x Zea luxurians | B73lux_BC1F2 | PRJNA639705 | SRR12023746 | NA | single | seedling | this study |
| B73lux_BC1F2_2 | Zea mays subsp.<br>mays x Zea luxurians | B73lux_BC1F2 | PRJNA639705 | SRR12023745 | NA | single | seedling | this study |
| B73lux_BC1F2_6 | Zea mays subsp.<br>mays x Zea luxurians | B73lux_BC1F2 | PRJNA639705 | SRR12023744 | NA | single | seedling | this study |
| B73lux_BC1F2_11 | Zea mays subsp.<br>mays x Zea luxurians | B73lux_BC1F2 | PRJNA639705 | SRR12023743 | NA | single | seedling | this study |
| B73lux_BC1F2_13 | Zea mays subsp.<br>mays x Zea luxurians | B73lux_BC1F2 | PRJNA639705 | SRR12023742 | NA | single | seedling | this study |
| B73lux_BC1F2_15 | Zea mays subsp.<br>mays x Zea luxurians | B73lux_BC1F2 | PRJNA639705 | SRR12023741 | NA | single | seedling | this study |
| B73lux_BC1F2_16 | Zea mays subsp.<br>mays x Zea luxurians | B73lux_BC1F2 | PRJNA639705 | SRR12023740 | NA | single | seedling | this study |
| B73lux_BC1F2_17 | Zea mays subsp.<br>mays x Zea luxurians | B73lux_BC1F2 | PRJNA639705 | SRR12023739 | NA | single | seedling | this study |
| B73lux_BC1F2_19 | Zea mays subsp.<br>mays x Zea luxurians | B73lux_BC1F2 | PRJNA639705 | SRR12023738 | NA | single | seedling | this study |
| trangenic_WT_1 | Zea mays | mixed | PRJNA639705 | SRR12023737 | NA | single | seedling | this study |
| trangenic_WT_2 | Zea mays | mixed | PRJNA639705 | SRR12023735 | NA | single | seedling | this study |
| trangenic_WT_3 | Zea mays | mixed | PRJNA639705 | SRR12023734 | NA | single | seedling | this study |
| trangenic_Ox_1 | Zea mays | mixed | PRJNA639705 | SRR12023733 | NA | single | seedling | this study |
| trangenic_Ox_2 | Zea mays | mixed | PRJNA639705 | SRR12023732 | NA | single | seedling | this study |
| trangenic_Ox_3 | Zea mays | mixed | PRJNA639705 | SRR12023731 | NA | single | seedling | this study |
| MS71 | Zea mays | MS71 | PRJNA305893 | SRR2994641 | NA | paired | leaves |  |
| Mo18W | Zea mays | Mo18W | PRJNA305893 | SRR3018349 | NA | paired | leaves |  |
| B97 | Zea mays | B97 | PRJNA305893 | SRR3018373 | NA | paired | leaves |  |
| CML333 | Zea mays | CML333 | PRJNA305893 | SRR3018392 | NA | paired | leaves |  |
| P39 | Zea mays | P39 | PRJNA305893 | SRR3018404 | NA | paired | leaves |  |
| Il14H | Zea mays | Il14H | PRJNA305893 | SRR3018410 | NA | paired | leaves |  |
| KI11 | Zea mays | KI11 | PRJNA305893 | SRR3018575 | NA | paired | leaves |  |
| Ky21 | Zea mays | Ky21 | PRJNA305893 | SRR3018597 | NA | paired | leaves |  |
| Tx303 | Zea mays | Tx303 | PRJNA305893 | SRR3018625 | NA | paired | leaves |  |
| NC358 | Zea mays | NC358 | PRJNA305893 | SRR3018741 | NA | paired | leaves |  |
| KI3 | Zea mays | KI3 | PRJNA305893 | SRR3018808 | NA | paired | leaves |  |
| Hp301 | Zea mays | Hp301 | PRJNA305893 | SRR3018811 | NA | paired | leaves |  |
| CML69 | Zea mays | CML69 | PRJNA305893 | SRR3018813 | NA | paired | leaves |  |
| CML247 | Zea mays | CML247 | PRJNA305893 | SRR3018814 | NA | paired | leaves |  |
| TZl8 | Zea mays | TZl8 | PRJNA305893 | SRR3018816 | NA | paired | leaves |  |
| M162W | Zea mays | M162W | PRJNA305893 | SRR3018819 | NA | paired | leaves |  |
| CML103 | Zea mays | CML103 | PRJNA305893 | SRR3018820 | NA | paired | leaves |  |
| OH7B | Zea mays | OH7B | PRJNA305893 | SRR3018821 | NA | paired | leaves |  |
| OH43 | Zea mays | OH43 | PRJNA305893 | SRR3018822 | NA | paired | leaves |  |
| CML322 | Zea mays | CML322 | PRJNA305893 | SRR3018825 | NA | paired | leaves |  |
| CML52 | Zea mays | CML52 | PRJNA305893 | SRR3018826 | NA | paired | leaves |  |
| CML228 | Zea mays | CML228 | PRJNA305893 | SRR3018827 | NA | paired | leaves |  |
| M37W | Zea mays | M37W | PRJNA305893 | SRR3018833 | NA | paired | leaves |  |
| B73_1 | Zea mays | B73 | SRP105290 | SRR5466739 | NA | single | seedling | Gent et al.,<br>2017 |
| B73_2 | Zea mays | B73 | SRP105290 | SRR5466555 | NA | single | seedling | Gent et al.,<br>2017 |
| B73_3 | Zea mays | B73 | SRP105290 | SRR5466556 | NA | single | seedling | Gent et al.,<br>2017 |
| B73_5 | Zea mays | B73 | SRP105290 | SRR5466737 | NA | single | seedling | Gent et al.,<br>2017 |
| B73_6 | Zea mays | B73 | SRP105290 | SRR5466390 | NA | single | seedling | Gent et al.,<br>2017 |
| J178-3 | Zea luxurians | PI 422162 | SRP105290 | SRR5466387 | SRR5466389 | single | seedling | Gent et al.,<br>2017 |
| Oax-2 | Zea mays mays | PI 628470 | SRP105290 | SRR5466588 | SRR5466710 | single | seedling | Gent et al.,<br>2017 |

